# Responsiveness variability during anaesthesia relates to inherent differences in brain structure and function of the fronto-parietal networks

**DOI:** 10.1101/2020.06.10.144394

**Authors:** Feng Deng, Nicola Taylor, Adrian M. Owen, Rhodri Cusack, Lorina Naci

## Abstract

Anaesthesia combined with functional neuroimaging provides a powerful approach for understanding the brain mechanisms of consciousness. Although propofol is used ubiquitously in clinical interventions that reversibly suppress consciousness, it shows large inter-individual variability, and the brain bases of this variability remain poorly understood. We asked whether three networks key to conscious cognition — the dorsal attention (DAN), executive control (ECN), and default mode (DMN) — underlie responsiveness variability under anaesthesia. Healthy participants (N=17) were moderately anaesthetized during narrative understanding and resting state conditions inside the Magnetic Resonance Imaging scanner. A target detection task measured behavioural responsiveness. An independent behavioural study (N=25) qualified the attention demands of narrative understanding. 30% of participants were unaffected in their response times, thus thwarting a key aim of anaesthesia — the suppression of behavioural responsiveness. Individuals with stronger functional connectivity within the DAN and ECN, between them, and to the DMN, and with larger grey matter volume in frontal regions were more resilient to anaesthesia. For the first time, we show that responsiveness variability during propofol anaesthesia relates to inherent differences in brain structure and function of the fronto-parietal networks, which can be predicted prior to sedation. Results highlight novel markers for improving awareness monitoring during clinical anaesthesia.

**Key points:**

- Moderate propofol anaesthesia showed highly variable effects across individuals
- 30% of healthy participants’ response times were unaffected by anaesthesia and 70% had significantly delayed, fragmented, or fully omitted responses
- Grey matter volume in frontal cortex and functional connectivity of the fronto-parietal networks predicted resilience to anaesthesia

## 1. Introduction

Understanding how the human brain gives rise to consciousness remains a grand challenge for modern neuroscience. A first step to understanding consciousness is to define what it is not. To this end, anaesthesia combined with functional neuroimaging provides a powerful approach for studying the brain mechanisms that change as consciousness fades (Vatansever et al, 2020; Pal et al., 2020; Varley et al., 2020; Luppi et al., 2019; Demertzi et al., 2019; Mashour and Hudetz 2018; Naci et al., 2018; MacDonald et al., 2015; Sarasso et al., 2015; Stamatakis et al., 2010), and, conversely, that are necessary for realizing human consciousness. Anaesthesia has been used for over 150 years to reversibly abolish consciousness in clinical medicine, but its effect can vary substantially between individuals. At moderate dosages, the intended suppression of behavioural responsiveness is highly variable (Chennu et al., 2016; Bola et al., 2019), and at deep anaesthesia dosages, in rare cases (0.1-0.2%; Mashour and Avidan 2015; Sandin et al., 2000), individuals retain conscious awareness, also known as ‘unintended intraoperative awareness’ (Pandit et al., 2017; Sanders et al., 2017; Sandin et a., 2000; Mashour and Avidan 2015). A much higher percentage of patients presumed to be unconscious during general anaesthesia (22%; Leslie et al., 2007) may have subjective experiences, such as dreaming. The brain bases of this considerable inter-individual effect variability remain poorly understood.

A key and mostly overlooked question is what these individual differences (Searle and Hopkins 2009; Palanca et al., 2009) can reveal about the unravelling of conscious cognition as consciousness falters during anaesthesia. Propofol is the most common anaesthetic agent in clinical interventions that require the reversible suppression of consciousness. Two recent electroencephalography (EEG) studies that measured variable behavioural responsiveness during mild and moderate propofol anaesthesia reported that the participants’ varying levels of responsiveness were differentiated by alpha band connectivity during wakefulness (Chennu et al., 2016), and changing patterns of EEG signal diversity from wakefulness to moderate sedation (Bola et al., 2019). However, the brain regional- or network-bases for this substantial individual variability (Searle and Hopkins 2009; Palanca et al., 2009), and its potential link to cognitive function, remain unknown. Propofol-induced sedation produces selective metabolic impairment and reduces brain activity bilaterally in frontal and parietal associative regions (Witon et al., 2020; Boly et al., 2008; Plourde et al., 2006; Alkire 2005; Baars et al., 2003; Fiset et al., 1999; for a review see MacDonald et al., 2015; but see Pal et al., 2020). Therefore, three brain networks with frontal and parietal lobe distribution, the dorsal attention network (DAN), executive control network (ECN), and default mode network (DMN) are primary candidate sources of individual variability under propofol sedation.

The DAN and ECN, distributed laterally across frontal and parietal lobes, are key to orchestrating stimulus-driven and goal-directed cognition (Duncan 2010; Elliott 2003; Corbetta and Schulman 2002; Kroger et al., 2002; Shallice 1988). The DMN extends partially in lateral and medial frontal and parietal lobes and is involved in internally-oriented cognition (Andrews-Hanna et al. 2010; Schneider et al., 2008; Buckner et al., 2008; Beer 2007; D’Argembeau et al., 2005; Wicker et al., 2003; Gusnard et al., 2001), but also external environment monitoring (Vatansever et al., 2017; Spreng et al. 2014; Buckner et al. 2008; Hahn et al., 2007) and shifts in contextually relevant information (Smith et al., 2018). The DAN, ECN and DMN are selectively impaired during loss of consciousness across conditions, e.g., under anaesthesia and after severe brain injury (Luppi et al., 2019; Naci et al., 2018).

Furthermore, the DAN/ECN and the DMN display an antagonistic relationship (Huang et al., 2020; Fox et al., 2005; Fransson et al., 2005) during spontaneous thought and goal-directed cognition. During conditions that engage external attention, the reduction of functional activity in the DMN (Greicius et al., 2003, Raichle et al., 2001, Shulman et al. 1997), is concomitant with an increase of activity in the DAN and ECN (Seeley et al. 2007; Dosenbach et al. 2007; Fox et al., 2005; Sridharan et al. 2008). The anticorrelation between these is related to individual differences in performance variability (Kelly 2008), and directly supports sustained attention (Kucyi et al., 2020). Conversely, this antagonistic relationship breaks down in state-related manipulations of consciousness, such as during sleep (Tagliazucchi et al., 2013), anaesthesia (Bonhomme at el., 2012; Boveroux et al., 2010), and severe brain injury (Haugg et al., 2018; Boly et al., 2009). Despite this accumulating evidence for the key and interrelated roles of the DAN, ECN, and DMN in supporting conscious cognition (Huang et al., 2020; Demertzi et al., 2013; Vanhaudenhuyse et al., 2011), and their primacy as target sites of propofol-induced sedation, their roles in individual differences under anaesthesia have not been previously investigated.

To address this gap, in two studies we tested whether variability or impairments in functional connectivity within and between the DAN, ECN, and DMN underlie individual differences in responsiveness during propofol anaesthesia. To directly investigate their individual and joint roles, in a first functional Magnetic Resonance Imaging (fMRI) study we scanned healthy participants (N=17) during wakefulness and during the administration of propofol at dosages of ‘moderate anaesthesia,’ expressly aimed at engendering individual differences. Variability in behavioural responsiveness in each state was assessed with an auditory target detection task prior to scanning. To test whether any individual differences in responsiveness were related to differences in perceptual or high-level attention processes that were invisible to the clinical sedation scale (Ramsay et al., 1974), participants were scanned during an active listening condition comprised of a brief (5 minutes) engaging auditory narrative, and during the condition of resting state. In a second behavioural study, we assessed the high-level attention demands of narrative understanding in an independent behavioural group (N=25) and related them to the brain activity of participants who underwent scanning.

## 2. Methods

### 2.1. Participants

*Study 1*. Healthy participants for the fMRI anaesthesia study (N=17; 18-40 years; 13 males) were tested in the MRI scanner at the Robarts Research Institute, Western University, in Canada. Ethical approval was obtained from the Health Sciences Research Ethics Board and Psychology Research Ethics Board of Western University. *Study 2.* An independent group of healthy participants (N=25; 18-40 years; 7 males) were tested behaviourally at the Global Brain health Institute at Trinity College Dublin, in Ireland. Ethical approval was obtained from the School of Psychology Research Ethics Board, Trinity College Dublin. All healthy participants were right-handed, native English speakers, had self-reported normal hearing and no history of neurological disorders. Informed consent was obtained for each participant prior to the experiment.

### 2.2. Stimuli and Design

#### Study 1

The study comprised a structural scan at the start, followed by two experimental sessions, one during wakefulness and the other during moderate anaesthesia, each with the same design. At the start of each session, a clinical assessment of sedation (Ramsay, 1974) was conducted to confirm wakefulness or moderate anaesthesia. Subsequently, a behavioural task, and two fMRI acquisitions, (a) a free listening to an auditory narrative (same in both sessions) and (b) a resting state scan, were conducted in each session. Listening to plot-driven narratives is naturally engaging, requires minimal behavioural collaboration from participants, and therefore, is highly suitable for testing perceptual or high-level attention processes independently of behavioural output or eye opening (Naci et al., 2018; 2017), which are impaired in moderate anaesthesia. Due to design constrains, the resting state scan followed the auditory target detection task and the narrative scan, and spontaneous thought in the resting state may have been influenced by these preceding activities. However, as this study’s primary question did not concern the resting state, this is not a substantive limitation. The data analysed here was collected as part of a larger study, and the wakefulness data has previously been reported in relation to deep anaesthesia (Naci et al. 2018). See below for full protocol details.

##### Behavioural testing

###### Auditory target detection task

Before commencing the scanning for either session, participants were asked to perform a computerized auditory target detection task (50 trials), which aimed to assess individual responsiveness differences during moderate anaesthesia. fMRI was not acquired during this task. Sound was presented with the Sensimetrics (S14; www.sens.com) headphones. A volume level deemed comfortable by each individual was determined prior to testing and used for the duration of the experiment. Participants were instructed to press a button with their index finger as soon as they heard an auditory beep (1200 Hz, 100 ms duration, 10 ms ramps) and to keep their eyes on the fixation cross on the screen. Participant had up to 3000 ms to make a response. Once a response was made, a pause of 1500 ms occurred prior to the presentation of the next stimulus. The stimulus onset asynchrony (SOA) = min [reaction time (RT), 3000ms] + 1500 ms. The SOA was jittered by virtue of variable RTs between trials. To ensure the participant did not lose contact with the response box during moderate anaesthesia, the response box was attached securely to their dominant hand. Individual and group RTs were used for behavioural analyses.

##### MRI testing

Inside the MRI scanner, participants underwent two functional scans during wakefulness and moderate anaesthesia. A plot-driven auditory narrative (5 minutes) was presented over MRI compatible noise cancelation headphones (Sensimetrics, S14; www.sens.com). Participants were asked to simply listen with eyes closed. The narrative comprised a highly engaging auditory excerpt from the movie ‘Taken’ depicting dramatic events, where a young girl travelling abroad without her family is kidnapped while speaking on the phone to her father. The same narrative was presented during wakefulness and moderate anaesthesia. A similar eyes-closed, resting state condition (8 minutes) was also acquired in each state for comparison with the narrative condition.

##### Sedation Procedure

The level of sedation was measured with the Ramsay clinical sedation scale for each participant by 3 independent assessors (two anaesthesiologists and one anaesthesia nurse) in person inside the scanner room. Before entering the fMRI scanner, a 20G I.V. cannula was inserted into a vein on the dorsum of the non-dominant hand of the participants. The propofol infusion system was connected to the cannula prior to the first scanning session. No propofol was administered during the wakeful session. Participants were fully wakeful, alert and communicated appropriately (Ramsay 1) and wakefulness (eye-opening) was monitored with an infrared camera placed inside the scanner. At the commencement of the moderate anaesthesia session, intravenous propofol was administered with a Baxter AS 50 (Singapore). An effect-site/plasma steering algorithm was used combined with the computer-controlled infusion pump to achieve step-wise increments in the sedative effect of Propofol. This infusion pump was manually adjusted to achieve the desired levels of sedation, guided by targeted concentrations of Propofol, as predicted by the TIVA Trainer (the European Society for Intravenous Anaesthesia, eurosiva.eu) pharmacokinetic simulation program. The pharmacokinetic model provided target-controlled infusion by adjusting infusion rates of Propofol over time to achieve and maintain the target blood concentrations as specified by the Marsh 3 compartment algorithm for each participant, as incorporated in the TIVA Trainer software (Marsh et al., 1991). In accordance with the Canadian Anaesthesia Society (CAS) guidelines, non-invasive blood pressure (NIBP), heart rate, oxygen saturation (SpO2) and end-tidal carbon dioxide (ETCO2) were monitored continuously through the use of a dedicated MR compatible anaesthesia monitor. Complete resuscitation equipment was present throughout the testing.

Propofol infusion commenced with a target effect-site concentration of 0.6 μg/ml and oxygen was titrated to maintain SpO2 above 96%. Throughout sedation, participants remained capable of spontaneous cardiovascular function and ventilation. Supplemental oxygen was administered via nasal cannulae to ensure adequate levels of oxygen at all times. If the Ramsay level was lower than 3 for moderate sedation, the concentration was slowly increased by increments of 0.3 *μ*g/ml with repeated assessments of responsiveness between increments to obtain a Ramsay score of 3. During administration of propofol, participants generally became calm and slowed in their response to verbal communication. Once participants stopped engaging in spontaneous conversation, and speech became sluggish, they were classified as being at Ramsey level 3. Nevertheless, when asked via loud verbal communication, participants agreed to perform the auditory target detection task as would be expected at Ramsey level 3. A brief recall task was adopted from the Mini Mental State Exam (Folstein et al., 1975), as a behavioural test complimentary to the anaesthesiologists’ assessments of whether sedation commensurate with Ramsay level 3 had been achieved. At the start of each session (wakefulness/moderate anaesthesia sessions), the researcher named three different sets of unrelated objects clearly and slowly and asked the participant to name each of them. The participant was instructed to remember the words in order to be able to repeat them in a short while. For 10 minutes following the word presentation, the participant was allowed to rest and performed no other task/was not exposed to any experimental stimuli. Subsequently, the participant was asked to repeat the words. Two different lists (a. Ball-Flag-Tree; b. Flower-Egg-Rope), counterbalanced across participants, were used to avoid familiarity effects between the wakefulness and moderate anaesthesia states. As expected, during the wakefulness session, all participants correctly repeated all three words, whereas during the moderate anaesthesia session, performance was varied with participants sluggishly repeating one or more of the words, consistent with Ramsay level 3 sedation. During moderate anaesthesia (Ramsey 3), the mean estimated effect-site propofol concentration, as provided by the TIVA Trainer software, was 1.99 (1.59-2.39) *μ*g/ml and the mean estimated plasma propofol concentration was 2.02 (1.56-2.48) *μ*g/ml.

#### Study 2

To provide a subjective measure of attention during the same story, an independent behavioural group of participants that did not undergo scanning, rated how “suspenseful” this story was every 2 seconds, from ‘least’ (1) to ‘most suspenseful’ (9). The audio excerpt was divided into 156 clips, each 2 seconds long to match the repetition time (TR, 2 sec) used in the independent participant group from Study 1 (who underwent MRI scanning at the Robarts Research Institute, at Western University, in Canada). Participants heard the stimuli through over-ear headphones connected to the stimulus presentation computer and used the keyboard to record responses, in a sound-isolated laboratory at the Global Brain Health Institute, at Trinity College Dublin, in Ireland. Participants had up to 3000 ms to make a response, at which point the next sequential clip began immediately. At the end of the experiment, participants indicated via a feedback questionnaire that the interruptions did not disrupt the coherence of the story’s plot and their perception of suspense throughout.

### 2.3. Data Analyses

#### Study 1

##### Analysis of behavioural data

As responses beyond 3000ms were not monitored, for the purposes of behavioural analyses, we considered participants who did not respond within 3000ms during anaesthesia as very delayed in their response time rather as missing data points. Participants were awake and responsive but very sluggish, consistent with moderate anaesthesia/Ramsay level 3 designation, and, therefore, it is very likely that responses were made outside the 3000ms window. Individual-level differences in reaction times (RTs) of the target detection task between wakefulness and moderate anaesthesia sessions were assessed by independent samples t-tests. The rate of RT change from baseline was calculated as follows: [(RT during anaesthesia - RT during wakefulness)/RT during wakefulness]*100. The hit rate was calculated as the number of responses divided by the number of trials for wakefulness and moderate anaesthesia separately. The Mahalanobis distance was computed to detect outliers in our multivariate dataset (Hadi 1992), comprised of the rate of RT change from baseline and hit rate during anaesthesia. Participants were considered outliers if their robust Mahalanobis distance from the rest of the distribution was significant at p < 0.01.

##### MRI Data Acquisition

Functional images were obtained on a 3T Siemens Prisma system, with a 32-channel head coil. The high-resolution brain structural images were acquired using a T1-weighted 3D MPRAGE sequence with the following parameters, voxel size: 1 × 1 × 1 mm, TA=5 minutes and 38 seconds, echo time (TE) = 4.25ms, matrix size=240×256×192, flip angle (FA) = 9 degrees. The functional echo-planar images (EPI) images were obtained with following parameters, 33 slices, voxel size: 3 × 3 × 3, inter-slice gap of 25%, repetition time (TR) = 2000ms, TE=30ms, matrix size=64×64, FA=75 degrees. The audio narrative and resting state had 155 and 256 scans, respectively.

###### A) Functional data preprocessing

Standard preprocessing procedures and data analyses were performed with SPM8 (Statistical Parametric Mapping; Wellcome Institute of Cognitive Neurology, http://www.fil.ion.ucl.ac.uk/spm/software/spm8/) and the AA pipeline software (Cusack et al., 2015). In the preprocessing pipeline, we performed slice timing correction, motion correction, registration to structural images, normalization to a template brain, and smoothing. The data were smoothed with a Gaussian smoothing kernel of 10mm FWHM (Peigneux et al. 2006). Spatial normalization was performed using SPM8’s segment-and-normalize procedure, whereby the T1 structural was segmented into grey and white matter and normalized to a segmented MNI-152 template. These normalization parameters were then applied to all EPIs. The time series in each voxel was high pass-filtered with a cutoff of 1/128 Hz to remove low-frequency noise, and scaled to a grand mean of 100 across voxels and scans in each session. Prior to analyses, the first five scans of each session were discarded to achieve T1 equilibrium and to allow participants to adjust to the noise of the scanner. To avoid the formation of artificial anti-correlations (Murphy et al. 2009; Anderson et al., 2011), we performed no global signal regression.

###### B) Structural data preprocessing

The brain structural images were processed using FreeSurfer package (https://surfer.nmr.mgh.harvard.edu/), a well-documented automated program which is widely used to perform surface-based morphometric analysis (Dale et al. 1999). The processing steps include: 1) removing the non-brain tissue; 2) transforming the skull-stripping brain volume to Talairach-like space; 3) segmenting brain tissues to GM, WM, cerebrospinal fluid (CSF); 4) performing intensity normalization to remove the effect of bias field; 5) building a surface tessellation to generate a triangular cortical mesh consisting of about 300,000 vertices in the whole brain surface; 6) correcting topological deficits of cortical surface; and 7) deforming brain surface to generate optimized models of GM/WM and GM/CSF boundaries.

##### Analyses of fMRI data

Group-level correlational analyses explored, for each voxel, the cross-subject synchronization of brain activity by measuring the correlation of each participant’s time-course with the mean time-course of all other participants (Naci et al., 2018; 2017; 2014). Further, to investigate whether the behavioural variability under anaesthesia related to perceptual or higher-order processing differences among participants, we performed two mixed data-driven and model-based analyses. First, we extracted the sound envelope of the auditory narrative via the Matlab *MIRtoolbox* (http://www.jyu.fi/hum/laitokset/musiikki/en/research/coe/materials/mirtoolbox) and built a generalised linear model (GLM) by using statistical parametric mapping to derive auditory characteristic-related brain activation for each individual. Subsequently, the z-scored average suspense ratings of the narrative obtained by the independent group of participants in study 2 (see SI) were used as a regressor in the fMRI data of individuals who underwent propofol anaesthesia in study 1, to measure the neural correlates of perceptual or high-level attention processes during the narrative condition. The regressors were generated by convolving boxcar functions with the canonical hemodynamic response function (Friston et al., 1989). For the task activation analyses, head movement was accounted for by regressing out the 6 motion parameters at the individual level. Included in the general linear model were nuisance variables, comprising the movement parameters in the three directions of motion and three degrees of rotation, as well as a constant (all-ones vector) that served to regress out the session mean. (Additional analyses of motion showed no significant differences between participants groups. See Table S4). Fixed-effect analyses were performed in each subject, corrected for temporal auto-correlation using an AR (1)+white noise model. Linear contrasts were used to obtain subject-specific estimates for each effect of interest. Linear contrast coefficients for each participant were entered into the second level random-effects analysis. Clusters or voxels that survived at p<0.05 threshold, corrected for multiple comparisons with the family-wise error (FWE) were considered statistically significant. For thresholding at the cluster level, an uncorrected cluster forming voxel threshold of p<0.001 was used.

To test whether individual differences under anaesthesia related to inherent brain features independent from the propofol sedation, we investigated the functional connectivity between and within the three networks during wakefulness, and in sedation. Functional connectivity (FC) within and between the DAN, ECN, and DMN was assessed by computing the Pearson correlation of the fMRI time courses between 19 regions of interests (ROIs; spherical, 10mm diameter) (from Raichle 2011) constituting these different brain networks, as identified by resting state studies (Table 1). This parcellation method is theoretically-driven based on meta-analyses of these three particular networks, and, furthermore, helps to relate our current findings to our previous findings based on the same parcellation method (Naci et al., 2018; Haugg et al., 2018). Permutation (1000 times) tests were used to explore FC difference between conditions and were false discovery rate (FDR) corrected for multiple comparisons. Independent analyses considered the contribution of signals from the white matter and cerebrospinal fluid to functional connectivity (Table S3). All of the analyses were conducted using Fisher z-transformed correlation (Pearson *r*). Glass’s delta was used to compute the effect size of the comparison between wakeful and moderate anaesthesia states, because of the different standard deviations of the two states, while Hedges’ *g* was used to compute the effect size of the comparison between fast and slow participants because of the different sample sizes for the two groups.

**Table 1.** Overview of selected regions of interests for the three functional networks

| Network/Region | ROI | MNI coordinates |  |  |
| --- | --- | --- | --- | --- |
|  |  | x | y | z |
| **Default Mode Network** | Posterior cingulate/precuneus | 0 | -52 | 27 |
|  | Medial prefrontal | -1 | 54 | 27 |
|  | Left lateral parietal | -46 | -66 | 30 |
|  | Right lateral parietal | 49 | -63 | 33 |
|  | Left inferior temporal | -61 | -24 | -9 |
|  | Right inferior temporal | 58 | -24 | -9 |
| **Dorsal Attention Network** | Left frontal eye field | -29 | -9 | 54 |
|  | Right frontal eye field | 29 | -9 | 54 |
|  | Left posterior IPS | -26 | -66 | 48 |
|  | Right posterior IPS | 26 | -66 | 48 |
|  | Left anterior IPS | -44 | -39 | 45 |
|  | Right anterior IPS | 41 | -39 | 45 |
|  | Left MT | -50 | -66 | -6 |
|  | Right MT | 53 | -63 | -6 |
| **Executive Control Network** | Dorsal medial PFC | 0 | 24 | 46 |
|  | Left anterior PFC | -44 | 45 | 0 |
|  | Right anterior PFC | 44 | 45 | 0 |
|  | Left superior parietal | -50 | -51 | 45 |
|  | Right superior parietal | 50 | -51 | 45 |

Abbreviations: IPS: Intraparietal sulcus, MT: Middle temporal area, PFC: Prefrontal cortex

##### Grey matter volume analyses

To test whether individual differences under anaesthesia related to inherent brain features independent from the propofol sedation, we also tested grey matter volume differences across participants. Vertex-wise grey matter volume (GMV) was computed (Winkler et al., 2018) and smoothed with a 10-mm full-width at half maximum Gaussian kernel to perform the statistics. Monte Carlo simulation cluster analysis and a cluster-wise threshold of p <0.05 was adopted for multiple comparisons correction (Hagler et al., 2006). Then, averaged GMV values for each significant cluster, for each individual participant, was extracted to perform permutation tests (1000 times) that tested for clusters/regions showing significant GMV differences between fast and slow participants. The FDR method was used to correct for multiple comparisons.

#### Study 2

To determine how similar suspense ratings were across the group, the inter-subject correlation of suspense ratings was computed as the average of the Pearson correlations of each participant’s data with the mean data from the rest of the group To account for the non-normalized distribution of correlation values (Fisher 1915), all statistical analyses were performed on z-transformed correlation values, using Fisher’s r-to-z transformation. For visualization purposes, we re-transformed these z-values in correlation values.

## 3. Results

### 3.1 The effect of moderate anaesthesia on behavioural responsiveness

Despite the same Ramsay 3 clinical score, independently determined by three assessors, we observed significant heterogeneity in the RTs of the auditory detection task (Figure 1A). 5/17 participants were not delayed significantly relative to their wakeful responses, 9/17 were significantly delayed and had fragmented responses (showing 2–40% missing trials), and 3/17 failed to make any responses within the 3000ms time window (Figure 1A–B), despite agreeing to have understood the task instructions and to make responses via the button box affixed to their dominant hand. This large individual variability was at odds with the propofol infusion rates titrated for each participant based on the pharmacokinetic model adjusted for demographic variables (Marsh et al., 1991), to maintain stable target blood concentrations consistent with Ramsay 3 sedation level (Ramsay, 1974).

**Figure 1.**
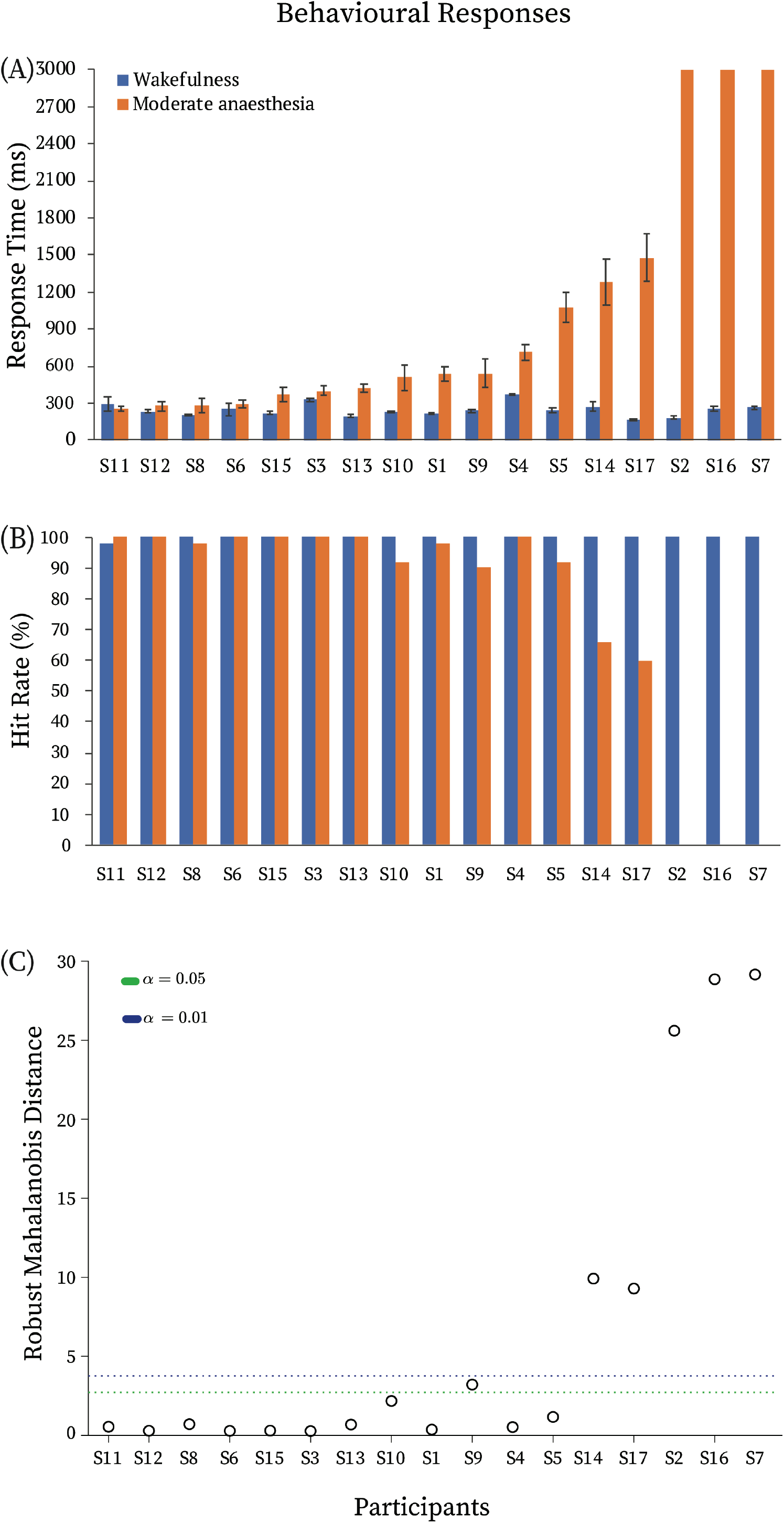
Behavioural responses. (A) Reaction times and (B) hit rates during the auditory target detection task in the wakeful (blue) and moderate anaesthesia (orange) states. (C) Multivariate outlier detection based on the robust Mahalanobis distance measure. Green/blue dotted lines depict the significance threshold of p<0.05/p<0.01. For further analyses, participants that showed robust Mahalanobis distance significantly different from the rest at p<0.01 (above the blue line) were labelled ‘slow’ participants (N=5), whereas the rest (below the blue line) were labelled ‘fast’ participants (N=12). The horizontal axis represents each individual (S1-S17=Subject1-Subject17).

Based on the rate of RT change from the wakeful baseline and hit rates during anaesthesia (Figure 1B), multivariate data outlier detection showed that 5 participants had significantly different robust Mahalanobis distance from the rest (Figure 1C). On this basis, individuals were divided into two groups for further analyses: ‘fast’ participants (FPs; N=12), and ‘slow’ participants (SPs; N=5). Individual responsiveness variability was not explained by demographic differences, as suggested by no differences in age (between-samples t-test, t = 1.3, p = 0.2) and gender (Fisher’s exact test, odds ratio = 1.3, p = 1) between FPs and SPs.

Subsequently, we tested whether the brain bases of this variability was related to three other factors. (1) Individual variability could be related to underlying differences in perceptual or high-level attention processes, which may have been invisible to the behavioural examination during the Ramsay assessment. (2) Responsiveness differences may be due to inherent connectivity differences prior to sedation, and/or alterations of connectivity due to sedation, within and between the DAN, ECN and DMN networks. (3) Individual responsiveness differences may be related to inherent structural brain differences, which may, in turn, also manifest as differences in functional connectivity. In the following analyses, we explored each of these factors in turn.

### 3.2 Perceptual and high-level attention processes and behavioural responsiveness during moderate anaesthesia

(1) We investigated whether each individual’s sensory-driven auditory perceptual processes, and high-level attention processes during the narrative condition related to their response times during the target detection task. First, we used SPM to model the relationship between the narrative’s perceptual properties captured by the sound envelope, and changes in brain activity over time. Then, each individual’s brain activity was related to their RTs in the target detection task. The sound envelope predicted brain activity in bilateral auditory cortex and right inferior frontal gyrus during wakefulness (Figure 2A), which as expected was dramatically reduced in moderate anaesthesia, to one small subthreshold cluster in left auditory cortex (p=0.05 FWE corrected) (Figure 2B). We observed no correlation between the participants’ reaction times in either wakefulness or moderate anaesthesia and the extent of their individual activations in auditory regions (Figure 2E–F). Furthermore, there were no significant differences in brain activation between the fast and slow groups for the processing of the sound envelope.

**Figure 2.**
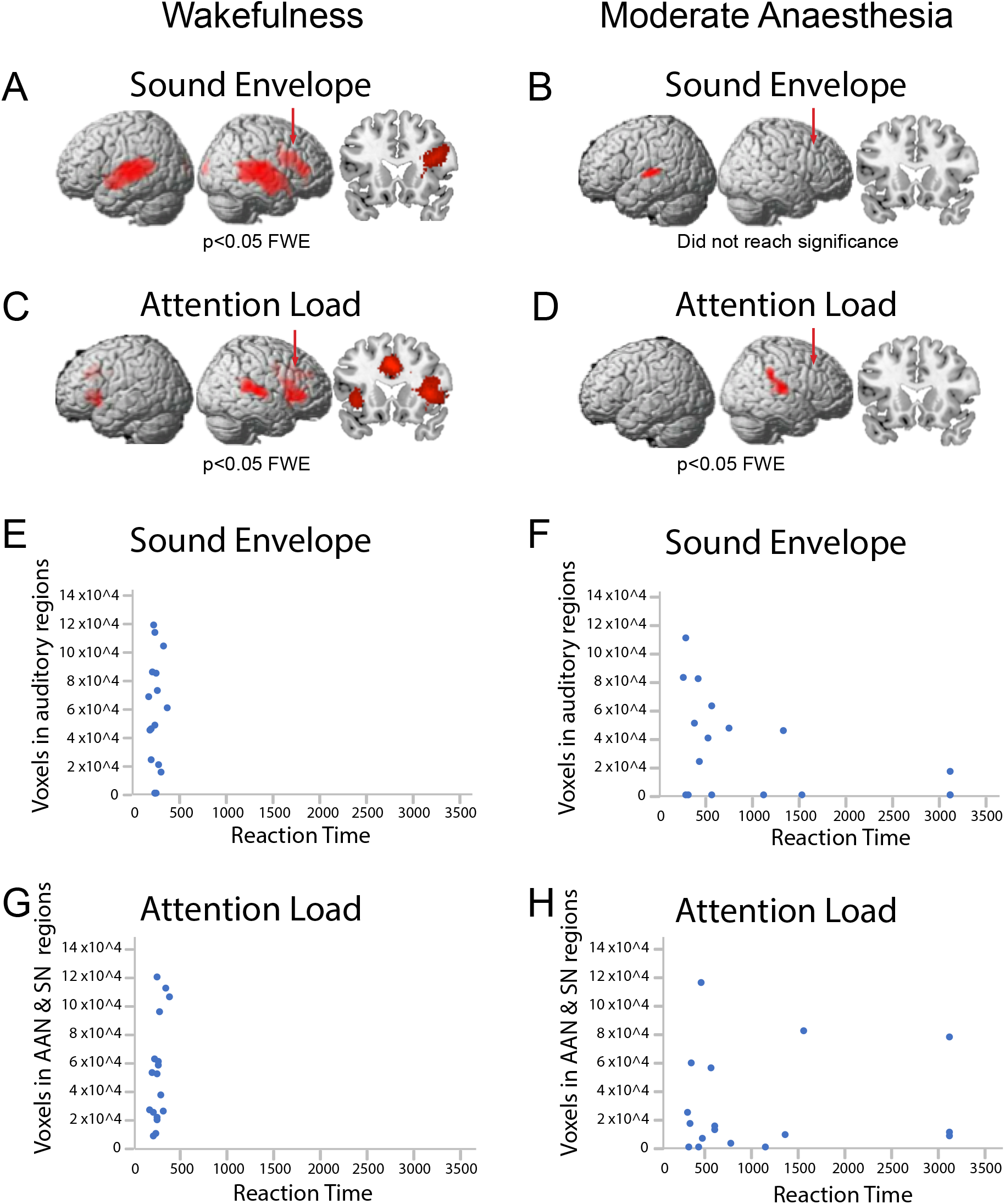
Perceptual and high-level attention processes during the narrative. At the group level, the sound envelope of the narrative predicted significant (p<0.05; FWE corrected) clusters in the (A) auditory cortex and right inferior frontal gyrus during wakefulness, and (B) left auditory cortex in moderate anaesthesia [did not reach statistical significance (p=0.05 FWE corrected)]. (C)/(D) At the group level, the suspense ratings of the audio narrative predicted significant (p<0.05; FWE corrected) clusters in the (C) auditory attention and salience networks during wakefulness, and (D) auditory attention network during moderate anaesthesia. Red arrows indicate the location of the coronal views with respect to the anterior-posterior dimension. (E)–(H) Panels display the number of voxels in primary auditory regions, auditory attention regions, and salience networks of individual participants that were predicted by the sound envelope and suspense ratings, relative to each individual’s reaction time in the target detection task. At the individual level, the sound envelope predicted significant activations in the auditory regions in (E) 14/17 of participants during wakefulness, and (F) 10/17 of participants during moderate anaesthesia. At the individual level, the suspense ratings predicted significant activations in the auditory attention and salience network regions in (G) 17/17 of participants during the wakefulness and (H) 14/17 of participants during moderate anaesthesia. There was no correlation between the perceptual or higher-order processes during the narrative and reaction times, during the target detection task, either in wakefulness (E)/(G) or in moderate anaesthesia (F)/(H). Abbreviations: AAN: Auditory Attention Network; SN: Salience Network.

Second, to investigate differential recruitment of high-level attention processes during the narrative, we used a previously established qualitative measure of the listeners’ sustained attention throughout the narrative — the perception of suspense on a moment-by-moment basis (Naci et al., 2014; 2017). We measured the perception of suspense in Study 2, in an independent group from those who underwent scanning. Suspense ratings throughout the narrative were highly and significantly correlated across individuals (r=0.90; SE=0.07) [t(24)=20.56, p<0.0001], confirming the similar high-level attention processes, and understanding of the narrative across different individuals. We used SPM to model the relationship between this qualitative measure of the narrative’s ongoing high-level attention demands and changes in brain activity over time, during the wakeful and moderate anaesthesia states. At the group level, during wakefulness, suspense ratings significantly (p<0.05; FWE corrected) predicted activity in regions of the auditory attention network (Tobyne et al., 2018; Michalka et al., 2015; Naci et al., 2013) and the salience network (Seeley 2019; Menon 2015) (Figure 2C), with clips rated as ‘highly suspenseful’ predicting stronger activity in these networks (Table 2). By contrast, during moderate anaesthesia, one small activation cluster was observed (Figure 2D).

**Table 2.** Coordinates for activation clusters associated with the perception of suspense during the narrative.

| Cluster Index | Cluster Peak Location | Peak MNI |  |  | Cluster-wise p-value |
| --- | --- | --- | --- | --- | --- |
|  |  | x | y | z |  |
| 1 | Cingulate Sulcus | -2 | 20 | 36 | <0.0001 |
| 2 | Right Superior Temporal Gyrus | 58 | -36 | 14 | <0.0001 |
| 3 | Right Anterior Insular Gyrus | 40 | 24 | 8 | <0.0001 |
| 4 | Posterior Cingulate Gyrus | 14 | -44 | 14 | <0.0001 |
| 5 | Left Anterior Insular Gyrus | -36 | 14 | -8 | <0.0001 |
| 6 | Thalamus | 4 | -20 | 10 | <0.005 |

As previously established (Naci et al. 2014; 2017; 2018), the similarity of the narrative’s suspense ratings enabled model-based predictions of the underlying brain activity, from the group data, in a leave-one-out fashion to individual participants. Using this approach, we related the activity in suspense-driven activation clusters for each individual to their respective RTs during the target detection task. No correlation was observed between the participants’ activation in the auditory attention and salience networks regions to their RTs (Figure 2 G–H). Furthermore, there were no significant differences in brain activation between the fast and slow groups for the processing of the narrative suspense.

### 3.3 Network connectivity and behavioural responsiveness during moderate anaesthesia

(2) We tested whether alterations in the connectivity within and between the DAN, ECN, and DMN during moderate anaesthesia (Figure S1, Supplemental Information) was associated with these individual differences (Figure 3). In the narrative condition, during wakefulness, functional connectivity within the DAN *(p* < 0.005), within the ECN (*p* < 0.005), between the DMN and DAN *(p* < 0.05), between DMN and ECN (*p* < 0.05), and between DAN and ECN (*p* < 0.05) differentiated the two response groups, with significantly higher FC for FPs relative to SPs (Figure 3B). During moderate anaesthesia, functional connectivity within the DMN (*p* < 0.05), within the ECN *(p* < 0.05), and between DAN and ECN (*p* < 0.05) differentiated FPs from SPs, with significantly higher FC for FPs relative to SPs (Figure 3B). In the resting state condition, during wakefulness, no differences were observed between FP and SP in any of connectivity types. During moderate anaesthesia, functional connectivity within and between the DAN, ECN, and DMN differentiated the two groups (DMN, *p* < 0.05; DAN, *p* < 0.05; ECN, *p* < 0.05; DMN-DAN, *p* < 0.005; DMN-ECN, *p* < 0.005; DAN-ECN, *p* < 0.005), with significantly higher FC for FPs relative to SPs (Figure 3B). All the above were permutation tests FDR corrected. These results were supported by analyses that considered the whole participant group (N=17). We found that the extent of RT change from wakefulness to sedation was significantly associated with functional connectivity (a) within the ECN, DMN–DAN and DMN–ECN in the wakeful narrative condition, and (b) within the ECN, DMN–DAN, DMN–ECN, DAN–ECN in the moderate sedation resting state (Table S1).

**Figure 3.**
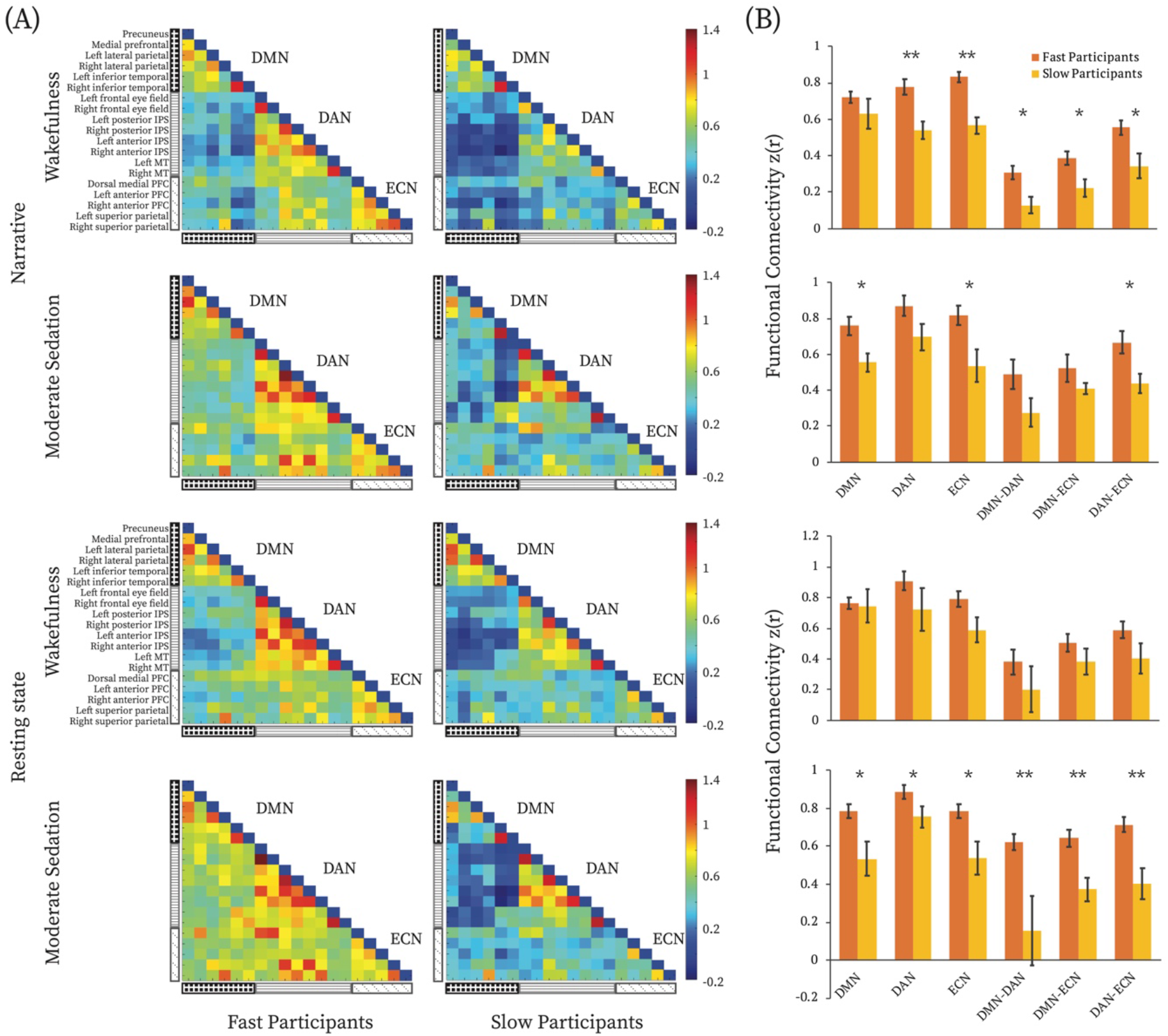
Functional connectivity differences between fast and slow participants. (A) FC matrices for the fast and slow participants during wakeful and moderate anaesthesia states in narrative and resting state conditions separately. The three black and white patterned rectangles on the left hand side represent, from top to bottom, the DMN, DAN and ECN. The colour bar indicates z-transformed Pearson correlation strength (low-high: blue-red). Each cell in the matrix represents FC between each node of the three networks. (B) Comparisons of averaged FC within and between the three brain networks, between fast and slow participants for each condition and state separately. * = p<0.05 FDR corrected for multiple comparisons. Abbreviations: DMN, default mode network; DAN, dorsal attention network; ECN, executive control network.

A 2×2×2 ANOVA explored the effects of *condition* (narrative, resting state), *state* (wakefulness, moderate anaesthesia) and *response group* (FPs, SPs), and their interactions. We found a main effect of response group (F (1,15) = 19.1, p < 0.005) that was driven by higher connectivity for FPs, and no effects on condition or state. However, effect size analyses showed higher difference between FPs and SPs in the narrative condition during wakefulness (Hedges’ g = 3.1) over all other conditions, including relative to moderate anaesthesia (Hedges’ g = 1.1), and relative to the resting state condition during wakefulness (Hedges’ g = 0.9), and moderate anaesthesia (Hedges’ g = 2.3).

### 3.4. Structural brain differences and behavioural responsiveness during moderate anaesthesia

(3) Next, we tested whether individual differences in behavioural responsiveness were related to structural brain differences, which may also underlie functional connectivity differences. To this end, we performed whole-brain vertex-based comparisons of grey matter volume between FPs and SPs. We found that FPs had uniformly significantly higher grey matter volume relative to the SPs, in two frontal regions, the left superior and dorsolateral frontal cortex (L SFC), including the presupplementary motor area (pre-SMA), and the left rostral middle frontal cortex (L rMFC) (Figure 4A, Table 3). The averaged GMV for each of these two regions were higher for FPs relative to SPs (L SFC, *p* < 0.0005, L rMFC, *p* < 0.0005; FDR corrected, Figure 4B). The opposite effect, i.e., higher grey matter for SP than FP, was not observed anywhere in the brain. The L SFC and L rMFC overlapped with the functional ROIs that defined the frontal nodes of the ECN (Table 1). Furthermore, when considering the whole participant group (N=17), we found that lower RT change from wakefulness to sedation was significantly associated with higher grey matter volume in the L SFC and the L rMFC (Figure S3, Table S2). These results further supported the functional connectivity results above (Figure 3).

**Figure 4.**
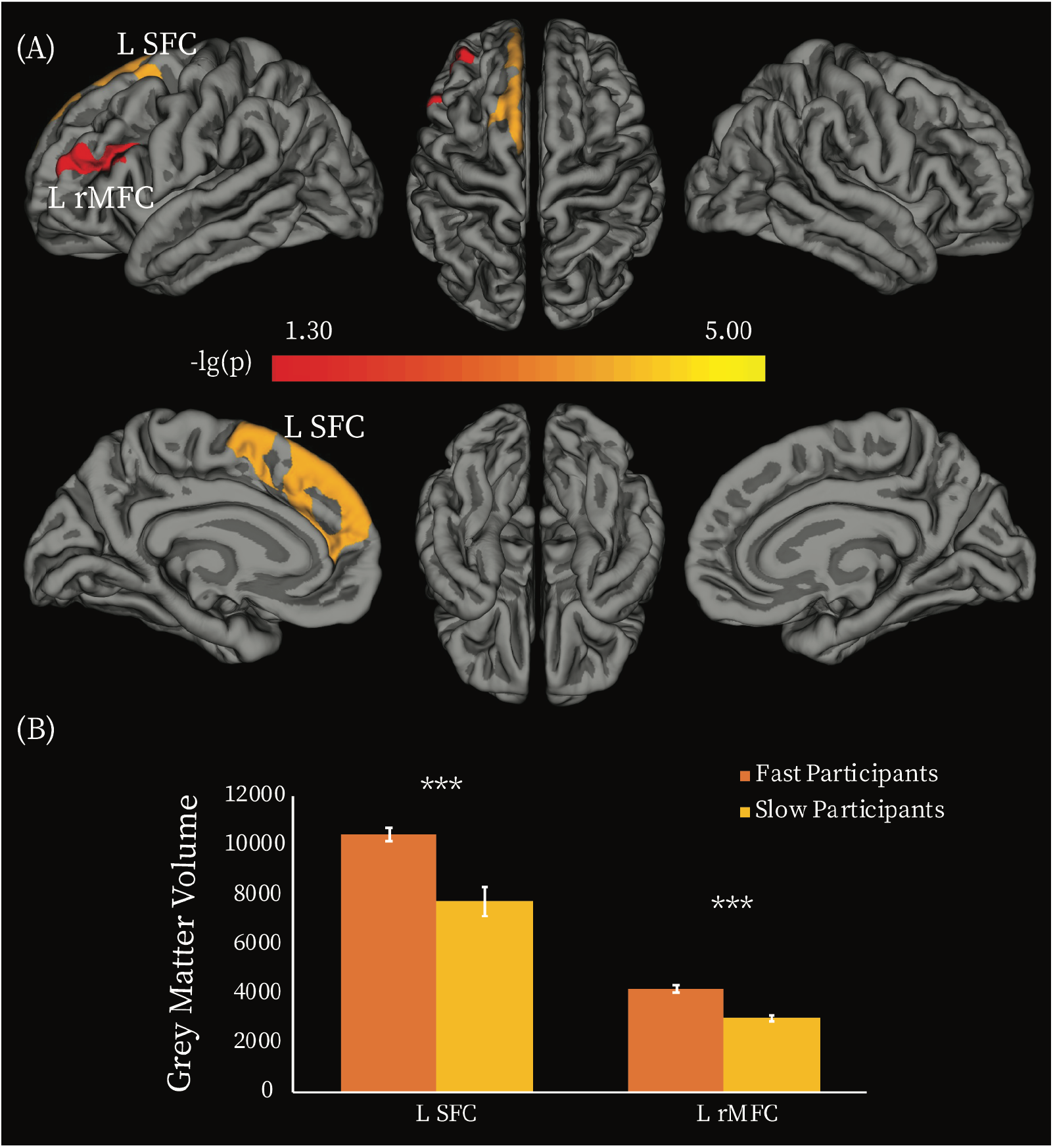
Grey matter volume differences between fast and slow participants. (A) Vertex-wise analysis. Two clusters showed significant between-group difference in GMV (p < 0.05, Monte Carlo simulation corrected). Colour bar indicates the p-value (-lg(p)). (B) Averaged GMV was extracted for each of the two clusters, and between-group comparisons of the averaged GMV were performed by using permutation tests. *** = p<0.0005 FDR corrected for multiple comparisons. Abbreviations: L SFC, left superior frontal cortex; L rMFC, left rostral middle frontal cortex.

**Table 3.**
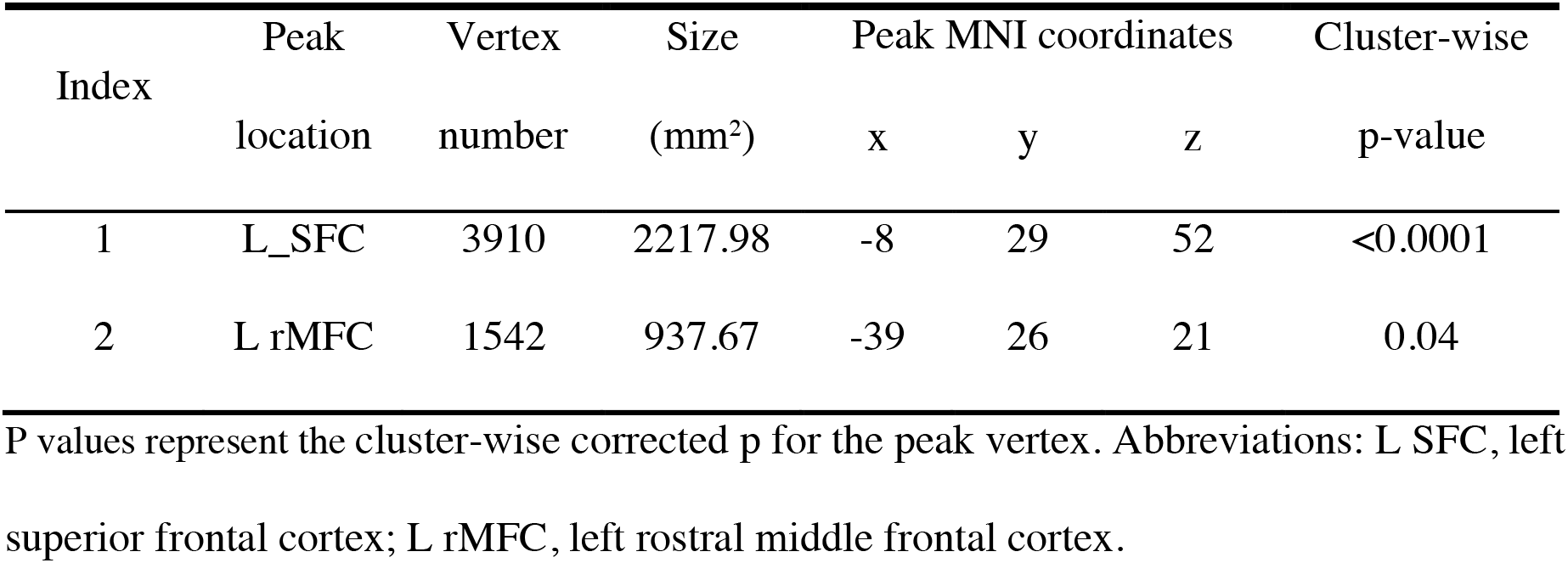
Regions showing significant grey matter volume difference between fast and slow participants.

We further tested whether the frontal and parietal aspects of the ECN were differentially involved in individual differences, by a direct comparison of the FC differences between FPs and SPs in the ROIs covering the frontal (dorsal medial PFC, left and right anterior PFC) and parietal (left and right superior parietal cortex) aspects of the ECN separately. A 2×2×2 ANOVA with factors *state* (wakefulness, moderate anaesthesia), *region* (frontal ECN, parietal ECN), and *response group* (FPs, SPs) showed a main effect of region [*F* (1,15) = 11.9, *p* < 0.005], driven by higher connectivity in the parietal aspect of the ECN, and a main effect of group [*F* (1,15) = 8.2, *p* < 0.05)], driven by higher connectivity in FPs relative to SPs. Direction comparisons between FPs and SPs in frontal and parietal aspects of ECN, showed that FC within the frontal, but not parietal, aspect of the ECN distinguished FPs and SPs during wakefulness and during moderate anaesthesia *(p* < 0.05, FDR corrected) (Figure 5).

**Figure 5.**
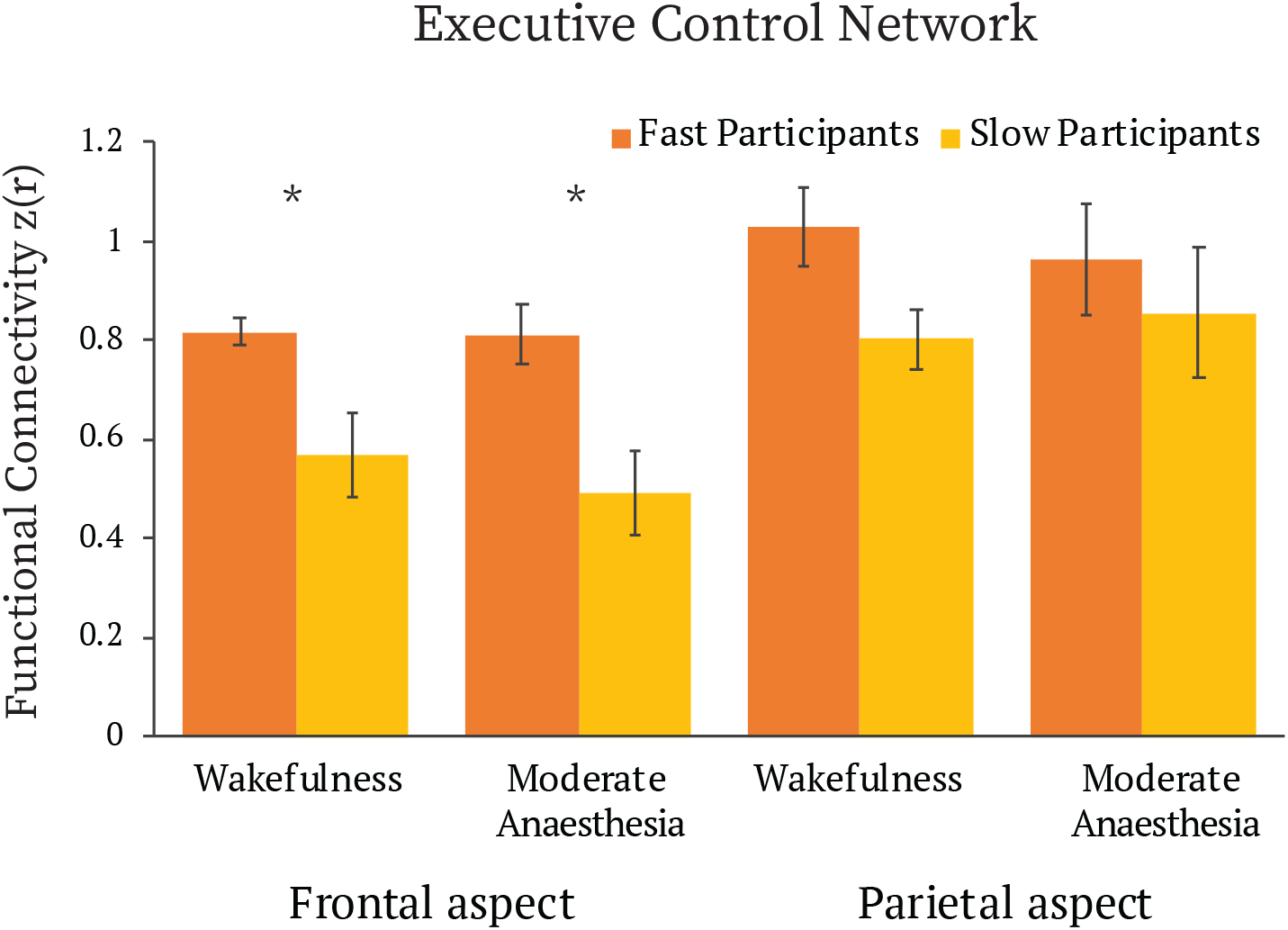
Differences of functional connectivity in frontal and parietal aspects of executive control network between fast and slow participants, in wakeful and moderate anaesthesia states in narrative condition. * = p <0.05 FDR corrected for multiple comparisons.

## 4. Discussion

Although anaesthesia has been used for over 150 years to reversibly abolish consciousness in clinical medicine, the brain bases of its considerable inter-individual effect variability remain poorly understood. To address this gap, we asked whether the connectivity within and between three networks that are primary sites of propofol-induced sedation and key to conscious cognition — the DAN, ECN and DMN — underlie individual responsiveness differences under anaesthesia. Consistent with previous studies (Chennu et al., 2016; Bola et al., 2019), we observed substantial individual differences in reaction times and hit rates under moderate anaesthesia. These were not related to differences in perceptual or high-level attention processes. Rather, the inherent level of functional connectivity during the narrative condition prior to sedation, within the DAN and ECN, between them, and to the DMN differentiated the participants’ responsiveness level. Furthermore, grey matter volume in the frontal cortex aspect of the ECN was significantly related to behavioural responsiveness during sedation. Participants whose reaction times showed little change from the wakeful baseline had significantly higher grey matter volume in these regions than participants who showed big changes. Therefore, for the first time, we show that responsiveness variability during propofol anaesthesia relates to differences in inherent brain structural and functional features of the fronto-parietal networks, which can be predicted prior to sedation.

Additionally, we found that during moderate anaesthesia, the connectivity within and between the DAN, ECN and DMN differentiated fast and slow participants, suggesting that perturbations by anaesthesia further exacerbate inherent inter-individual functional connectivity differences.

The differential impact of moderate anaesthesia on cognitive process can be inferred in a post hoc manner, based on the perceptual and cognitive processes recruited by the target detection task and the brain networks and regions that showed functional and/or structural differences between the participants. The task involved a hierarchy of perceptual and cognitive processes. These included (a) auditory perception, (b) understanding of linguistic instruction, (c) short-term memory to remember instructions for the duration of the experiment, (d) attention to auditory stimuli, (e) attention to intention to make a response, (f) cognitive control over other, competing and distracting mental processes, and the (g) execution of a motor response, when prompted by the auditory stimuli. The clinical assessment determined similar basic language communication (b), and short-term memory function (c) across participants during moderate anaesthesia. Furthermore, our results showed no relationship between behavioural responsiveness and sensory/auditory (a), or higher-order, including linguistic processes that relied on sustained attention (b, d), across participants. Therefore, differences in responsiveness were likely underpinned by differences in complex mental faculties, such as attention to intention (e), cognitive control (f) and action execution (g).

Consistent with this inference, our results demonstrated that FC within and between the DAN and ECN differentiated the participants’ level of responsiveness, suggesting that moderate anaesthesia impacts attention to intention and goal and action execution, subserved by the DAN and ECN (Marek and Dosenbach 2018; Dosenbach et al., 2007; Ernst and Paulus 2005). Furthermore, our structural imaging results demonstrated that grey matter volume differences in the frontal cortex aspect of the ECN, including the DLPFC and pre-SMA, underscore individual responsiveness differences. In addition to their role in action selection and execution (Lee et al., 1999; Boly et al., 2007), the DLPFC and pre-SMA represent higher-order attention, i.e., attention to intention. Lau et al (2004) investigated the brain regions that were preferentially involved in attending to the intention to move, relative to the actual movement, and found that brain activity in pre-SMA, as well as its functional connectivity with right DPFC represented the intention to move. The pre-SMA has also been implicated in sustained cognitive control (Nachev et al., 2005), another process that is key to successful task performance. Therefore, a post-hoc interpretation of our findings is that moderate anaesthesia impacts attention to intention, sustained cognitive control, and action execution, and that individuals with smaller grey matter volume in frontal regions and weaker functional connectivity within and between the DAN and ECN, may have a vulnerability for stronger suppression of behavioural responsiveness than those with higher values for these features. Additionally, the finding DMN connectivity during moderate anaesthesia differentiated fast and slow participants suggests that individual differences in responsiveness may also reflect the effect of sedation on self-referential processes (Andrews-Hanna et al. 2010; Schneider et al., 2008; Buckner et al., 2008; Beer 2007; D’Argembeau et al., 2005; Wicker et al., 2003; Gusnard et al., 2001), and context processing (Smith et al., 2018; Vatansever et al., 2017), which are supported by the DMN.

As the suppression of behavioural responsiveness is a key aim of anaesthesia, the converse finding was also highly relevant. Surprisingly, we found that 30% of participants showed no delay in reaction times under moderate anaesthesia relative to wakefulness, and critically, these exhibited inherent brain differences to participants who were significantly delayed. This result is highly relevant to the prediction of responsiveness variability under anaesthesia (Chennu et al., 2016; Palanca et al 2009) and monitoring depth-of-anaesthesia for the detection of unintended intraoperative awareness. Although rare (0.1-0.2%, Mashour and Avidan 2015; Sandin et al., 2000), unintended awareness can be very traumatic and lead to negative long-term health outcomes, such as post-traumatic stress disorder (up to 70%), as well as clinical depression or phobias (Pandit et al., 2017; 2014; Mashour and Avidan 2015). Given its rarity, unintended intraoperative awareness does not lend itself to direct investigation in the relatively small groups of typical research studies with vulnerable populations, including the present study, where individuals were anaesthetized without intubation in a research MRI setting.

Our results suggest that individuals with larger grey matter volume in frontal regions and stronger functional connectivity within DAN and ECN, between them and to the DMN, may require higher doses of propofol to become non-responsive to the same extent as individuals with weaker connectivity in these networks and smaller grey matter volume in frontal cortex. If replicated in a clinical context, these findings will provide novel markers that may help to improve the accuracy of awareness monitoring during clinical anaesthesia. We note that future clinical studies ought to measure the clearance of anaesthetic agents by each individual directly for, potentially, a more precise assessment of the metabolic variation across participants, than that provided by the anaesthesia delivery instrument used here. It is also worth noting that propofol was used here due to its wide prominence in clinical interventions, and future studies that employ the same paradigm across different agents will determine whether the brain-behaviour relationships we report here generalize to other anaesthetic agents.

The detection of individual responsiveness to anaesthesia prior to the sedation procedure is a question of clinical importance. We found that the wakeful narrative condition yielded the highest power relative to the other conditions for the detection of individual sedation responsiveness. This may be due to higher signal to noise during the narrative condition. Previous studies have shown that the active attention engagement during well-crafted narratives leads to reduced movement (Centeno et al., 2016) and less sleep in scanner (Centeno et al., 2016, Vanderwal et al., 2015), resulting in higher signal-to-noise relative to resting state (rs) fMRI (Wang et al., 2017). Moreover, it may be due to higher power to detect functional roles of brain networks during naturalistic stimuli relative to rsfMRI. Rs fMRI is entirely unconstrained, and thus it is difficult to separate signal due to cognitive processes from extraneous sources, including motion, cardiac and respiratory physiological noise, arterial CO2 concentration, blood pressure, and vasomotion (Murphy et al., 2013; Birn et al., 2006). By contrast, as we have previously established, naturalistic narratives drive diverging responses across brain networks, highlighting their functionally distinct roles, and thus may be more sensitive for investigating how their differing cognitive roles are impacted by anaesthesia than rsfMRI (Naci et al., 2018; Haugg et al., 2018; Naci et al., 2014). Our results are consistent with a growing recognition of the importance of naturalistic stimuli for studying cognitive processes and understanding the neural basis of real-world functioning (Sonkusare et al., 2019).

### 4.1 Methodological considerations

It is worth noting that this investigation included a relatively small sample size, not atypical of research studies with vulnerable populations, for aforementioned participant safety considerations. Future studies that will replicate findings in larger clinical groups are needed to further corroborate the present findings. Furthermore, as MRI technology improves, newer scanning parameters including shorter repetition time (TR) and higher spatial resolution could better capture spatiotemporal differences that underlie individual differences in response to anaesthesia.

## Data Availability Statement

In study 1, reaction time and fMRI data were collected in 2015, under ethics from Western University, Canada. This stipulated that data would be shared only with the research team directly involved in this study. Therefore, individual data cannot be freely shared due to ethics protocol regulation. However, we have uploaded group maps to NeuroVault (https://identifiers.org/neurovault.collection: 9576).

In study 2, suspense ratings data were collected under ethics from Trinity College Dublin, Ireland, which stipulated that anonymized data can be shared with the scientific community. Therefore, the behavioural data and code will be shared upon contacting the authors. The sharing practices are in line with the guidelines from the funding body.

