## Supporting Information File for "Responsiveness variability during anaesthesia relates to inherent differences in brain structure and function of the fronto-parietal networks"

**Title**

**Running title:** Brain bases of responsiveness variability

**Authors**

Feng Deng^a^, Nicola Taylor^a^, Adrian M. Owen^bd^, Rhodri Cusack^a^, Lorina Naci^ac*^

**Affiliations**

^a^ Trinity College Institute of Neuroscience, School of Psychology, Trinity College Dublin, Dublin, Ireland.

^b^Brain and Mind Institute, Western University, London, Canada.

^c^Global Brain Health Institute, Trinity College Dublin, Dublin, Ireland

^d^ Department of Physiology and Pharmacology and Department of Psychology, Western University, London, Canada.

^*^**Corresponding author:**

Lorina Naci

School of Psychology

Trinity College Institute of Neuroscience

Global Brain Health Institute

Trinity College Dublin

Dublin, Ireland

**Results**

### The effect of moderate anaesthesia on brain network connectivity across conditions

We investigated how moderate propofol anaesthesia perturbed the functional connectivity (FC) patterns relative to wakefulness across all the participants (Figure S1). FC matrices in wakeful and moderate anaesthesia states for the narrative and resting state conditions are shown in Figure S1A. For the narrative condition, a 2x6 ANOVA that explored the main effects of *state* (wakefulness, moderate anaesthesia) and *connectivity type* (DMN, DAN, ECN, DMN–DAN, DMN–ECN, DAN–ECN) showed significant main effects of state (F (1, 16) = 4.6, *p* < 0.05), connectivity type (F (3.1, 49.1) = 53.6, *p* < 0.0001), and a significant interaction effect of state by connectivity type (F (5, 80) = 7, *p* < 0.0001). We investigated further this interaction effect by testing the effect of anaesthesia state on each network connectivity. We found significantly higher DMN–DAN (permutation test, *p* < 0.05, FDR corrected), as well as DMN–ECN (permutation test, *p* < 0.05, FDR corrected) connectivity in moderate anaesthesia relative to wakefulness (Figure S1B). For the resting state condition, a similar 2x6 ANOVA showed a significant main effect of connectivity type (F (2.3, 36.7) = 37.3, *p* < 0.0001) and a significant interaction effect of state by connectivity type (F (2.8, 44.3) = 4.2, *p* < 0.05). We found no significant differences between wakefulness and moderate anaesthesia for each connectivity type (Figure S1B).

A direct comparison between the narrative and resting state conditions, with a 2x6 ANOVA [*condition* (narrative, resting state) x *connectivity type* (6 levels)] on the FC difference values (moderate anaesthesia - wakefulness), showed a significant main effect of connectivity type (F (5, 80) = 8.7, *p* < 0.0001), and no main effect of condition or interaction effect. However, effect size analyses showed higher effect sizes of moderate anaesthesia in the narrative relative to the resting state condition for both DMN–DAN (Glass’ Delta = 1.20 vs 0.52) and DMN–ECN (Glass’ Delta = 1.02 vs 0.5). In summary, these results suggested that the antagonistic relationship between the DMN and the DAN/ECN was reduced during moderate anaesthesia, with a stronger and significant result in the narrative condition relative to the resting state.

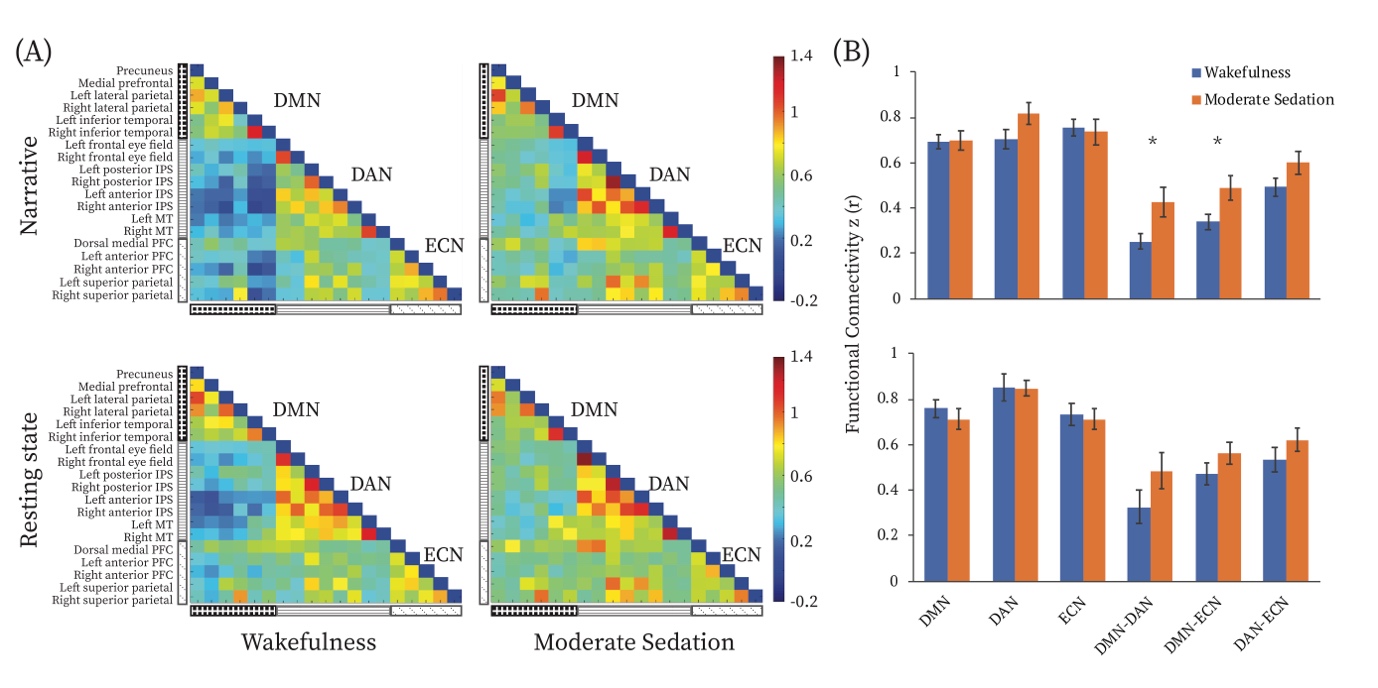

**Figure S1**. Functional connectivity (FC) in wakeful and moderate anaesthesia states for the narrative and resting state conditions. (A) Group-averaged FC connectivity matrices during wakeful and moderate anaesthesia states in the narrative and resting state conditions across all participants. The three black and white patterned rectangles on the left hand side represent, from top to bottom, the DMN, DAN and ECN. The colour bar indicates connectivity (measured as z-transformed Pearson correlation; low-high, blue-red). Each cell in the matrix represents FC between each node of the three networks. (B) Comparisons of FC averaged across all nodes for each network, within and between the three brain networks, during the wakeful and moderate anaesthesia states, in the narrative and resting state conditions separately. * = p <0.05 FDR corrected for multiple comparison comparisons. Abbreviations: DMN, default mode network; DAN, dorsal attention network; ECN, executive control network; IPS: intra parietal sulcus; PFC, prefrontal cortex; MT middle temporal cortex.

### Overall information processing during moderate anaesthesia

We used a previously established method (Naci et al., 2018; 2017; 2014), whereby the extent of stimulus-driven cross-subject correlation provides a proxy measure of regional stimulus-driven information processing. During wakefulness, we observed widespread and significant (p<0.05; FWE corrected) cross-subject correlation between healthy participants, bilaterally in sensory-driven auditory cortex, visual cortex, as well as higher-order frontal and parietal cortical regions (Figure S2). By contrast, during moderate anaesthesia, significant (p<0.05; FWE corrected) cross-subject correlation was observed only in auditory cortex and three small clusters (the left post central gyrus, left paracentral lobule and right precentral gyrus) of premotor and motor cortex, that have been implicated in language comprehension (Hauk et al., 2004) (Figure S2B). As expected, and consistent with previous studies (Naci et al., 2018; Davis et al., 2007), these results suggested reduced information processing during moderate anaesthesia relative to wakefulness.

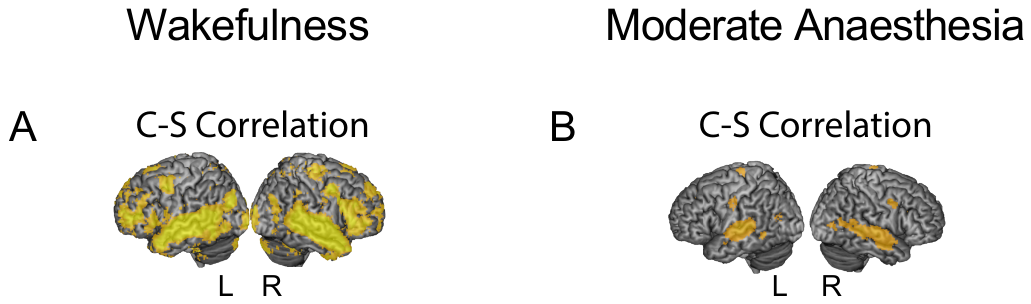

Figure S2. Effect of anaesthesia on information processing in the brain (A) At the group level, during wakefulness, the audio narrative elicited significant (p<0.05; FWE corrected) inter-subject correlation across the brain, including frontal and parietal cortex, thought to support high-level information integration over time. (B) At the group level, during moderate anaesthesia, significant (p<0.05; FWE corrected) inter-subject correlation was limited to the auditory cortex and three small clusters in regions of premotor and motor cortex that are involved in language comprehension.

### Associations between functional connectivity and behavioural responsiveness

In addition to the two group comparisons reported in the main text, we performed permutation tests to investigate the association of functional connectivity (FC) with the reaction time (RT) change from the awake state to moderate anaesthesia, in the whole cohort (N=17) (Table S1). In the narrative condition, during wakefulness, FC within the ECN (p < 0.05), between the DMN and DAN (p < 0.05), and between DMN and ECN (p < 0.05) were significantly negatively associated with the RT change. Higher FC was significantly associated with smaller changes in RT. These results were consistent with our main results obtained by comparing two groups, where we observed higher FC within the DAN (p < 0.005), within the ECN (p < 0.005), between the DMN and DAN (p < 0.05), between DMN and ECN (p < 0.05), and between DAN and ECN (p < 0.05) for fast participants (FPs; smaller RT change) relative to slow participants (SPs) in this condition (Figure 3B). During moderate anaesthesia of the narrative condition, FC was not significantly associated with the RT change. In the resting state condition, during wakefulness, FC was not significantly associated with the RT change. During moderate anaesthesia, FC within the ECN (p < 0.05), between the DMN and DAN (p < 0.005), between DMN and ECN (p < 0.05), and between DAN and ECN (p < 0.005) was significantly associated with the RT change. Higher FC was significantly associated with smaller RT change. This was consistent with our main results from the group comparisons showing that FC within and between the DAN, ECN, and DMN differentiated the two groups (DMN, p < 0.05; DAN, p < 0.05; ECN, p < 0.05; DMN-DAN, p < 0.005; DMN-ECN, p < 0.005; DAN-ECN, p < 0.005), with significantly higher FC for FPs (smaller RT change) relative to SPs in this condition (Figure 3B). All the above permutation tests were FDR corrected for multiple comparisons.

Table S1. Permutation tests for association of functional connectivity with the reaction time change from the awake state to the moderate anaesthesia state.

|  | Narrative | | | | Resting state | | | |
| --- | --- | --- | --- | --- | --- | --- | --- | --- |
|  | Wakefulness | | Moderate sedation | | Wakefulness | | Moderate sedation | |
|  | Coefficients | p values (Perm) | Coefficients | p values (Perm) | Coefficients | p values (Perm) | Coefficients | p values (Perm) |
| DMN | -175.9 | 0.69 | -875.8 | 0.28 | 466.5 | 0.71 | -925.0 | 0.20 |
| DAN | -1016.0 | 0.15 | -341.9 | 0.75 | -379.0 | 0.59 | -699.6 | 0.58 |
| ECN | -1692.0 | 0.03 | -1057.0 | 0.18 | -1108.0 | 0.58 | -1611.0 | 0.006 |
| DMN-DAN | -1678.0 | 0.046 | -569.7 | 0.28 | -300.6 | 0.71 | -1081.0 | 0.001 |
| DMN-ECN | -1875.0 | 0.03 | -389.8 | 0.42 | -602.9 | 0.59 | -1465.0 | 0.02 |
| DAN-ECN | -925.1 | 0.22 | -725.9 | 0.28 | -532.2 | 0.59 | -1580.0 | 0.004 |

Note: P values were obtained from permutation tests and were FDR corrected for multiple comparisons. Abbreviations: DMN, default mode network; DAN, dorsal attention network; ECN, executive control network; Perm, permutation tests.

### Associations between grey matter volume and behavioural responsiveness

In addition, we also analysed whether whole-brain grey matter volume was associated with the RT change in the entire participant group (N=17). We found significant associations between the RT change and grey matter volume, in two clusters in the left frontal cortex: the left superior and dorsolateral frontal cortex (L SFC) and the left rostral middle frontal cortex (L rMFC) (Figure S3, Table S2). Higher grey matter volume in these regions was significantly associated with smaller changes in RT under moderate anaesthesia relative to wakefulness. These results were consistent with our main results from the group comparisons, which showed that FPs (smaller RT change) had significantly higher grey matter volume relative to the SPs, in the L SFC and the L rMFC (Figure 4A, Table 3).

Table S2. Regions showing significant negative associations between the grey matter volume and the reaction time change from the awake state to the moderate anaesthesia state.

| Index | Peak location | Vertex number | Size (mm^2^) | Peak MNI coordinates | | | Cluster-wise p-value |
| --- | --- | --- | --- | --- | --- | --- | --- |
|  |  |  |  | x | y | z |  |
| 1 | L_SFC | 4730 | 2579.34 | -12 | -11 | 51 | 0.0003 |
| 2 | L rMFC | 3181 | 1997.57 | -38 | 25 | 26 | 0.002 |

Note: P values represent the cluster-wise corrected p for the peak vertex. Abbreviations: L SFC, left superior frontal cortex; L rMFC, left rostral middle frontal cortex.

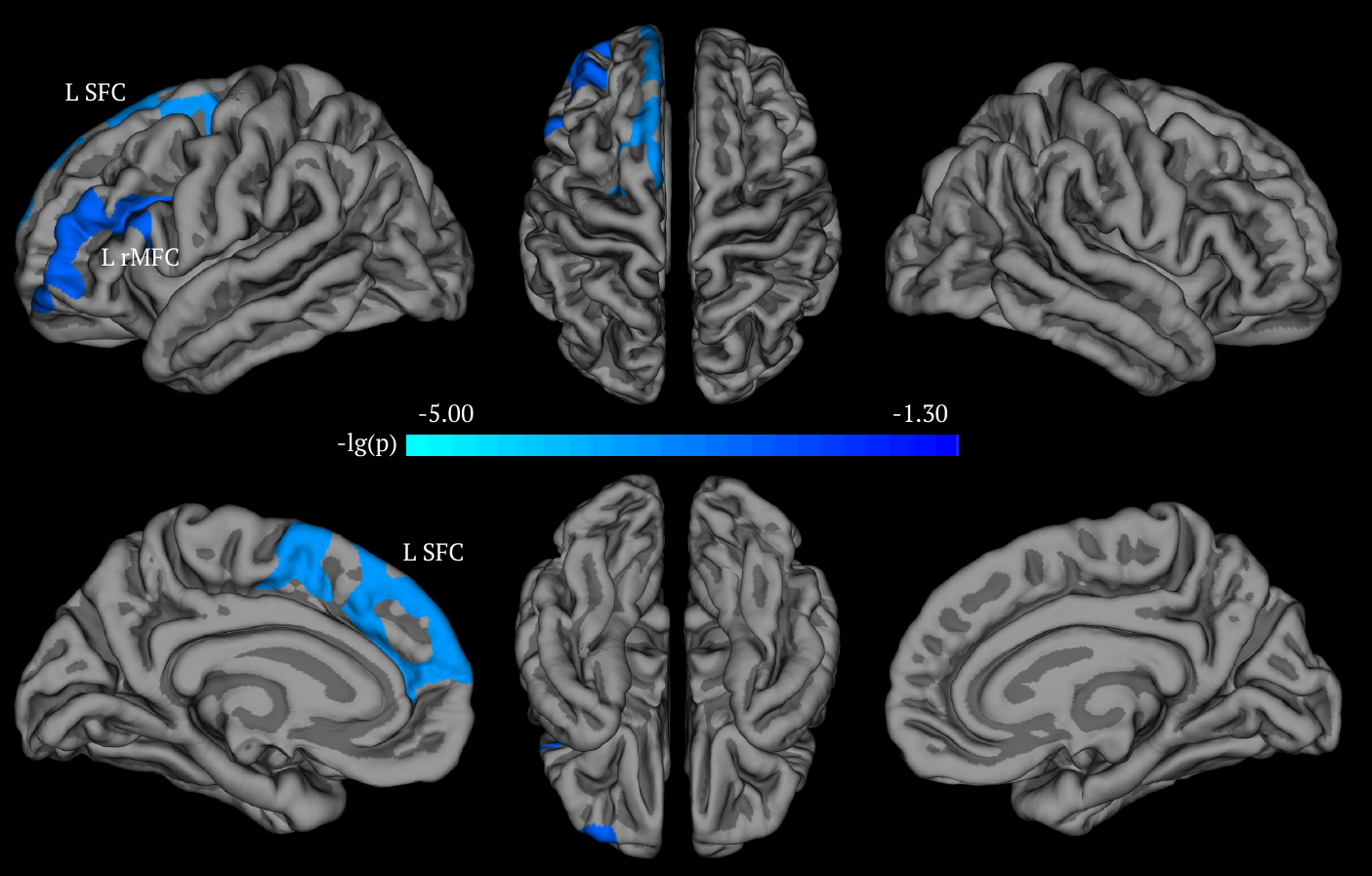

Figure S3. Vertex-wise analysis of the association between the grey matter volume and the reaction time change from the awake state to the moderate anaesthesia state. Two left frontal cortex clusters showed significant negative associations with the change RT (p < 0.05, Monte Carlo simulation corrected). The colour bar indicates the p-value (-lg(p)).

### Additional analyses accounting for white matter (WM) and cerebral spinal fluid (CSF) signals

To further account for the WM and CSF signals in functional connectivity, we performed additional analyses that controlled for these signals and included the 6 head motion parameters (Table S3). Consistent with the original results, in the narrative condition, during wakefulness, we found significantly lower FC within ECN (p = 0.002) in SPs relative to FPs.

Table S3. Permutation tests for group comparisons of functional connectivity between fast participants (FPs) and slow participants (SPs) after controlling for signals from the white matter (WM) and cerebrospinal fluid (CSF), and 6 head motion parameters at the individual level.

|  | Narrative | | | | Resting state | | | |
| --- | --- | --- | --- | --- | --- | --- | --- | --- |
|  | Wakefulness | | Moderate sedation | | Wakefulness | | Moderate sedation | |
|  | Group difference | p values (Perm) | Group difference | p values (Perm) | Group difference | p values (Perm) | Group difference | p values (Perm) |
| DMN | -0.03 | 0.35 | 0.03 | 0.40 | -0.06 | 0.25 | -0.05 | 0.40 |
| DAN | 0.08 | 0.21 | -0.01 | 0.40 | 0.10 | 0.22 | -0.13 | 0.06 |
| ECN | 0.20 | 0.002 | 0.15 | 0.19 | 0.08 | 0.22 | 0.02 | 0.40 |
| DMN-DAN | 0.11 | 0.21 | 0.02 | 0.40 | 0.10 | 0.25 | 0.14 | 0.24 |
| DMN-ECN | 0.08 | 0.22 | -0.07 | 0.40 | 0.03 | 0.35 | -0.05 | 0.32 |
| DAN-ECN | 0.06 | 0.24 | 0.02 | 0.40 | 0.08 | 0.22 | 0.02 | 0.40 |

Note: P values were obtained from permutation tests and reported after FDR correction for multiple comparisons. Abbreviations: DMN, default mode network; DAN, dorsal attention network; ECN, executive control network; Perm, permutation tests. Group difference = FPs-SPs

### Additional analyses accounting for head motion

For the functional connectivity analyses, we performed framewise displacement (FD) to evaluate the subjects with severe motion (more than 3 mm displacements and 3° rotations). In the narrative condition, no participant showed excessive head motion during wakefulness. 2 participants (S13 and S17) showed excessive head motion during moderate anaesthesia. In the resting state condition, 1 participant (S13) during wakefulness and 2 participants (S15 and S17) during moderate anaesthesia showed excessive head motion.

In the main text, we included all participants given no significant differences in the mean FD between FPs and SPs were observed in any conditions or states. To further account for the effect of the head motion, we performed permutation tests for the group comparisons with the mean FD regressed out and the main results stood (Table S4). Similarly, excluding participants with excessive head motion in specific conditions and states also did not change the main results (Table S4).

Table S4. Permutation tests for group comparisons of functional connectivity.

|  | Narrative | | | | Resting state | | | |
| --- | --- | --- | --- | --- | --- | --- | --- | --- |
|  | Wakefulness | | Moderate sedation | | Wakefulness | | Moderate sedation | |
|  | Group comparisons ^a^ | Group comparisons ^b^ | Group comparisons ^a^ | Group comparisons ^b^ | Group comparisons ^a^ | Group comparisons ^b^ | Group comparisons ^a^ | Group comparisons ^b^ |
| DMN | 0.11 | 0.11 | 0.03 | 0.02 | 0.37 | 0.38 | 0.008 | 0.01 |
| DAN | 0.004 | 0.004 | 0.04 | 0.07 | 0.17 | 0.17 | 0.04 | 0.04 |
| ECN | 0.004 | 0.004 | 0.03 | 0.01 | 0.10 | 0.05 | 0.009 | 0.01 |
| DMN-DAN | 0.004 | 0.004 | 0.02 | 0.048 | 0.21 | 0.21 | 0.002 | 0.002 |
| DMN-ECN | 0.01 | 0.01 | 0.08 | 0.14 | 0.21 | 0.21 | 0.003 | 0.002 |
| DAN-ECN | 0.01 | 0.01 | 0.02 | 0.02 | 0.17 | 0.17 | 0.003 | 0.002 |

Note: P values were obtained from permutation tests and reported after FDR correction for multiple comparisons. Abbreviations: DMN, default mode network; DAN, dorsal attention network; ECN, executive control network; Perm, permutation tests. Group comparison = FPs-SPs

^a^ permutation tests for group comparisons with the mean FD regressed out.

^b^ permutation tests for group comparisons after excluding participants with excessive head motion.
